# 2’3’-cGAMP triggers a STING and NF-κB dependent broad antiviral response in Drosophila

**DOI:** 10.1101/852319

**Authors:** Hua Cai, Andreas Holleufer, Bine Simonsen, Juliette Schneider, Aurélie Lemoine, Hans-Henrik Gad, Jingxian Huang, Jieqing Huang, Di Chen, Tao Peng, João T. Marques, Rune Hartmann, Nelson Martins, Jean-Luc Imler

## Abstract

We recently reported that an orthologue of STING regulates infection by picorna-like viruses in drosophila. In mammals, STING is activated by the cyclic dinucleotide 2’3’-cGAMP produced by cGAS, which acts as a receptor for cytosolic DNA. Here, we show that injection of flies with 2’3’-cGAMP can induce expression of dSTING-regulated genes. Co-injection of 2’3’-cGAMP with a panel of RNA or DNA viruses results in significant reduction of viral replication. This 2’3’-cGAMP-mediated protection is still observed in flies mutant for the genes *Atg7* and *AGO2*, which encode key components of the autophagy and small interfering RNA pathways, respectively. By contrast, it is abrogated in flies mutant for the NF-κB transcription factor Relish. Analysis of the transcriptome of 2’3’-cGAMP injected flies reveals a complex pattern of response, with early and late induced genes. Our results reveal that dSTING regulates an NF-κB-dependent antiviral program, which predates the emergence of Interferon Regulatory Factors and interferons in vertebrates.

## Introduction

Insects, like all animals, are plagued by viral infections, which they oppose through their innate immune system. Induction of transcription of antiviral genes upon sensing of infection is a common antiviral response observed across kingdoms. In insects, inducible responses contribute to defense against viruses, together with RNA interference (RNAi) and constitutively expressed restriction factors (reviewed in ^1^). Apart from RNAi, these mechanisms are still poorly characterized and appear to be largely virus-specific^2–4^. Combining genetics and transcriptomic analysis, we recently showed that the evolutionarily conserved factor dSTING participates together with the kinase IKKβ and the NF-κB transcription factor Relish in a novel pathway controlling infection by the picorna-like viruses Drosophila C virus (DCV) and Cricket Paralysis Virus (CrPV) in the model organism *Drosophila melanogaster*^5^.

In mammals, STING is a central component of the mammalian cytosolic DNA sensing pathway, where it acts downstream of the receptor cyclic GMP-AMP synthase (cGAS)^6^. Upon binding DNA, cGAS synthesizes 2’3’-cGAMP, a cyclic dinucleotide (CDN) secondary messenger that binds to and activates STING^7–10^. Bacteria also synthesize CDNs such as c-di-AMP, c-di-GMP and 3’3’-cGAMP, which can be sensed by STING (reviewed in ^11^). Upon activation, STING recruits through its C-terminal tail (CTT) region the kinase TBK1, which phosphorylates and activates the transcription factor Interferon Regulatory Factor (IRF) 3 to trigger interferon (IFN) production^12^. STING can also activate NF-κB and autophagy independently from its CTT domain in mammalian cells^13–15^.

The identification of STING in animals devoid of interferons, such as insects, raises the question of the ancestral function of this molecule. Since most invertebrates lack the IRF family of transcription factors (with the notable exception of mollusks) as well as interferons, the lack of a C-terminal tail (CTT) in invertebrate STING suggests that this recent addition co-evolved with the onset of IRF transcription factors and IFN cytokines^16^. In contrast, the ability of STING to regulate transcription factors of the NF-κB family^17 18 5^ or autophagy^19^, seems conserved throughout metazoa. Importantly, these responses are triggered in a CTT-independent manner in mammals^13–15^. Apart from the missing CTT, the global overall structure of STING is well conserved between vertebrates and invertebrates. Accordingly, *in vitro* studies with STING recombinant proteins from the sea anemone *Nematostella vectensis* (Cnidaria), the oyster *Crassostrea gigas* (Molluscs) and the worm *Capitella teleta* (Annelids) revealed that they all bind CDNs^20^. Intriguingly however, binding of CDNs was not observed with recombinant STING produced from several insect species, including drosophila^20^.

The mechanism by which STING exerts its antiviral effect in insects, which could provide important clues on its ancestral function, is still unclear. Here, we identify 2’3’-cGAMP as a potent agonist of dSTING *in vivo* and show that it triggers a strong Relish-dependent transcriptional response that confers protection against a broad range of RNA and DNA viruses.

## RESULTS

### A subset of CDNs trigger expression of STING dependent virus regulated genes

To characterize *in vivo* the dSTING pathway, we used *dSTING^Rxn^* (RXN) loss of function mutant flies (Fig.S1a). Expression of *dSTING* was significantly reduced by 9- to 27-fold in the mutant, as previously described, but was restored to wild type level when a genomic rescue was introduced in the flies (Fig.S1b). Basal levels or induction by DCV of three previously described IKKβ and dSTING dependent genes (CG13641, CG42825, and CG33926, hereafter referred to as *sting regulated gene* (*srg)1*, *srg2* and *srg3*, respectively) was significantly reduced in RXN mutant flies compared to *dSTING^Control^* (WT) or *dSTING^Rescue^* flies (Fig.S1c-e). By contrast, induction of the gene *Hsp26*^21^ by DCV was not affected in the RXN mutant (Fig.S1f). We noted that STING expression was still induced by DCV infection in RXN mutant flies, reaching levels close to wild type three days post infection (dpi, Fig.S1b).

We next analyzed whether the dSTING pathway could be activated by naturally occurring CDNs known to be agonists of STING in other organisms. CDNs injected into WT flies led to a significant increase in *dSTING* and *srg1-3* with c-di-AMP, 3’3’-cGAMP and 2’3’-cGAMP at 6 hours post injection (hpi, Fig.1a-d) and at 24 hpi (Fig.1e-h). Only c-di-GMP did not trigger a response in these experiments. The induction of *srg1-3* by CDNs was reduced in RXN mutant flies at 6 hpi or abolished at 24 hpi (Fig.1b-d and f-h). For *dSTING* itself, the pattern of induction was similar in RXN and WT flies, although the level of expression was always significantly reduced in the mutant flies (Fig.1a,e). Because neither the promoter nor the open reading frame (ORF) of the short form of dSTING are affected by the RXN deletion (Fig.S1a), we hypothesize that a residual level of dSTING protein in the mutant accounts for some remaining activity of the pathway. Induction of *srg1* and *srg2* by 2’3’-cGAMP was rapid, peaking at 3 or 6 hpi and decreasing afterwards (Fig.1i,j). Interestingly, inducible expression of *srg3* remained high at 24 hpi (Fig.1k). Induction of *srg1-3* by 2’3’-cGAMP was reduced or abolished in *Relish* null mutant flies (Fig.1l-n). Overall, these data reveal that a subset of naturally occurring CDNs can trigger gene expression in *Drosophila*, in a manner dependent on dSTING and Relish.

**Figure 1.**
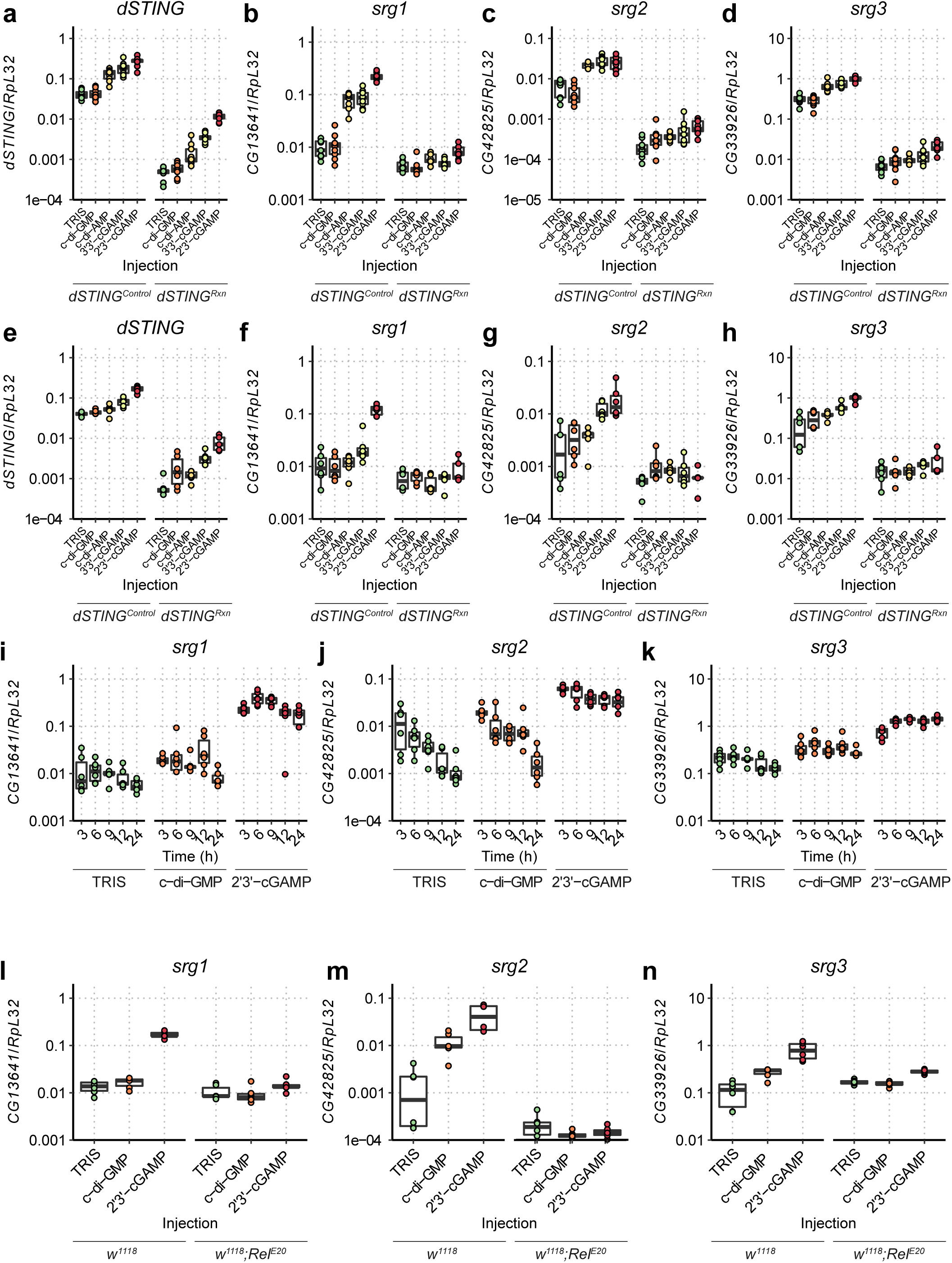
2’3’-cGAMP injection induces a dynamic dSTING-Relish dependent transcriptional response in *D. melanogaster*. (**a-h**) Relative gene expression of the indicated dSTING-regulated genes at 6h (**a-d**) and 24h (**e-h**) after injection of Tris and different CDNs in *dSTING^Control^* or *dSTING^Rxn^* mutant fies. *dSTING* and *srg1-3* were significantly induced in *dSTING^Control^* flies 6 hours post injection (hpi) with c-di-AMP, c-di-AMP, 3’3’-cGAMP and 2’3’-cGAMP (|*t*| ≥ 4.807, *p* < 0.001 for all comparisons of Tris vs CDN injections). c-di-GMP injection did not lead to significant changes in gene expression at any timepoint (|*t*| ≤ 0.184, *p* ≥ 0.184 for all comparisons of Tris vs c-di-GMP injected flies). srg1-3 were never significantly induced in *dSTING^Rxn^*mutants 24hpi (|*t*| ≤ 3.290, *p* ≥ 0.200 for all comparisons of Tris vs CDN injections). *dSTING* was induced in *dSTING^Rxn^* mutants (|*t*| > 2.963, *p* < 0.017, for all comparisons of Tris vs CDN injections, excluding c-di-GMP), but the level of expression was always significantly lower than in control flies (|*t*| > 19.043, *p* < 0.001 for all pairwise comparisons between control and *dSTING^Rxn^*). (**i-k**) Expression levels of *srg1-3* at different times post-injection with Tris, cyclic-di-GMP or 2’3’-cGAMP. (**l-n**) Expression levels of *srg1-3* 6h post-injection with Tris, cyclic-di-GMP or 2’3’-cGAMP in control *w^1118^* and *w^1118^;Rel^E20^*mutant flies. After 2’3’-cGAMP injection, induction folds of srg1-3 were always significantly lower in Relish mutant than in control flies (|*t*| > 5.48, *p* ≤ 0.001 for all comparison of differences in *srg1-3* levels between Tris and 2’3’-cGAMP injected flies). Data are from two independent experiments. Expression levels are shown relative to the housekeeping gene *RpL32* and are normalized by experiment.

### 2’3’-cGAMP has a significant impact on the transcriptome of whole flies

We next performed genome-wide transcriptomic analysis to identify 2’3’-cGAMP regulated genes in whole flies. We identified 427 up- and 545 down-regulated genes by more than 1.5-fold in animals injected with 2’3’-cGAMP compared to Tris buffer (Fig.2a), with 269, 88 and 115 transcripts upregulated and 311, 53 and 63 transcripts downregulated at the 6, 12 and 24h timepoints, respectively (Fig. S2). In contrast, only four up- and one down-regulated transcripts were observed when c-di-GMP was injected into WT flies (Table S1). Clustering analysis revealed three broad categories of up- and down-regulated genes based on their early, sustained and late kinetics of induction or repression (Fig.2a’, Table S2). Among upregulated genes, *srg1* was induced rapidly, while *srg3* classifies as a late induced gene, confirming our initial observation. Rapidly induced genes included antimicrobial peptides, cytokines such as *spaetzle* and *upd3*, transcription factors (e.g. *Rel*, *kay*, *Ets21C, FoxK*) and other signaling molecules (*Takl1*, *pirk*, *Charon*, *dSTING*) (Fig.2a’’). Interestingly, one of the three canonical components of the siRNA pathway, *AGO2*, was rapidly induced by 2’3’-cGAMP, together with *pst* and *ref(2)P*, which encode restriction factors against picorna-like viruses^22, 23^ and rhabdoviruses^24^, respectively. Late induced genes were mainly unknown but included the JAK-STAT regulated gene *vir-1* and the antiviral gene *Nazo* (Fig.2a’’)^5, 25^. Gene ontology analysis revealed that the early and sustained upregulated genes were significantly enriched for a single functional category, namely immunity (Fig.2b). No such enrichment was detected in the late induced genes. By contrast, the downregulated genes upon 2’3’-cGAMP injection were associated with mitochondria or belonged to several metabolic pathways, including carbohydrate, lipid and protein metabolism (Fig.2b). This points to an impact of CDN injection on metabolism, possibly reflecting cellular reprograming.

**Figure 2.**
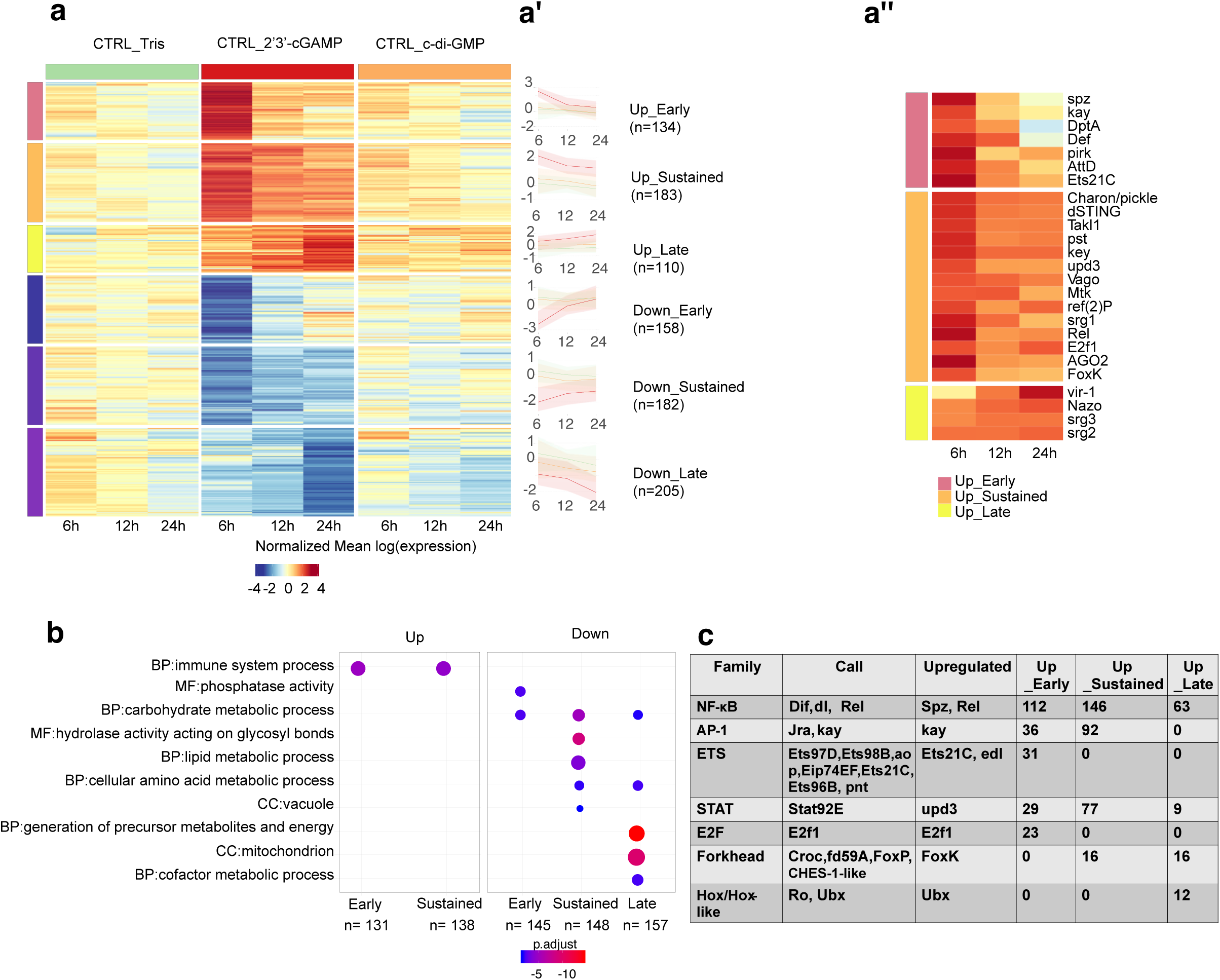
2’3’-cGAMP induces a strong transcriptional response in *D.melanogaster*. (**a**) Expression profiles of *dSTING^Control^* flies injected with Tris, 2’3’-cGAMP or c-di-GMP (6, 12 and 24h post-injection). All differentially expressed genes (DEG) between 2’,3’-cGAMP- and Tris-injected flies for at least one timepoint or on average across all time points are shown. Values are normalized to the mean log (expression) of Tris-injected flies across the three time points. Expression profiles of up- and down-regulated genes in 2’3’-cGAMP-injected flies were clustered by partition around medoids. (**a’**) Normalized mean gene expression by experimental condition in each temporal expression profiles and across the different timepoints. (**a’’**) Expression of some representative genes discussed in the text. (**b**) Gene ontology enrichment analysis of the DEGs across the different temporal expression profiles. **BP**: Biological process, **MF**: Molecular function, **CC**: Cellular compartment. Size and color of circles indicates respectively the number of DEG and p-value for the enrichment of each category. (**c**) Numbers of DEGs potentially regulated by **Upregulated** transcription factors and cytokines. Genes with high confidence binding sites for other TFs (**Call**) of the same family are included.

We next performed *in silico* analysis of predicted binding sites for transcription factors in the upregulated genes. We found that 75% of the upregulated genes (321) contained binding sites for members of the NF-κB family. While 84% of early and 80% of the sustained genes contained NF-κB binding sites, only 57% (63) of the late genes contained such binding sites, suggesting a significant secondary response at 24h post-cGAMP injection. Furthermore, we could find enrichment for binding sites for other upregulated transcription factors, or for transcription factors regulated by induced cytokines (e.g. upd3) (Fig. 2c). Among these, STAT appears to play an important role in all temporal expression profiles, with binding sites in 22%, 42% and 8% of the genes in the early, sustained and late categories. Others, such as Ets21c, E2F1 and AP1 may participate in the early phase of the response to 2’3’-cGAMP, given their enrichment only in the early and sustained up-regulated genes (Fig.2c).

### Injection of 2’3’-cGAMP protects flies against viral infections

We next addressed the functional consequences of activation of the dSTING pathway by CDN injection. Co-injection of 2’3’-cGAMP with DCV resulted in a significant decrease of viral RNA in WT flies. Such a protective effect of 2’3’-cGAMP was not observed in *RXN* mutant flies, indicating that it was dependent on dSTING (Fig.3a). Accordingly, 2’3’-cGAMP significantly improved the survival of DCV infected WT flies but not of *RXN* mutants (Fig.3b). Co-injection of 2’3’-cGAMP but not c-di-GMP also resulted in significantly reduced accumulation of viral RNA for four other RNA viruses, namely the positive strand RNA viruses cricket paralysis virus (CrPV) and Flock house virus (FHV), the negative strand RNA virus vesicular stomatitis virus (VSV), and the double strand DNA virus Kallithea virus (KV) (Fig.3c-f). Collectively, these results indicate that 2’3’-cGAMP triggers protection against a broad range of viruses.

**Figure 3.**
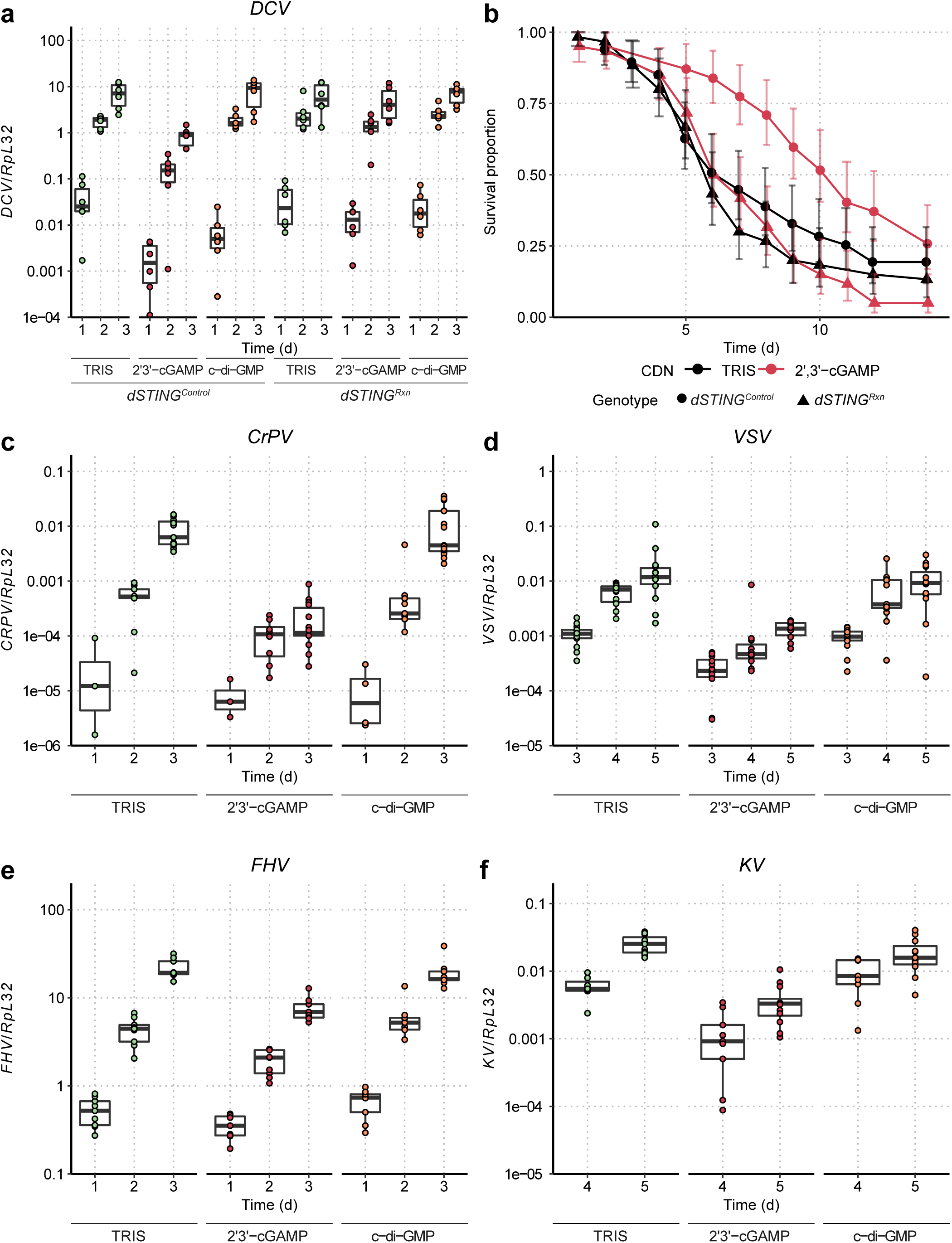
2’3’-cGAMP injection induces a broad, dSTING-dependent, antiviral protection in *D. melanogaster*. Relative DCV RNA loads (**a**) and survival after infection (**b**) of *dSTING^Control^*and *dSTING^Rxn^* mutant flies after co-injection of DCV and Tris, 2’3’-cGAMP or c-di-GMP at different days post-injection (d). Co-injection with 2’3’-cGAMP resulted in a significant decrease of viral RNA in *dSTING^Control^* flies 2 and 3 dpi (|*t*| ≥ 3.466, *p* ≤ 0.002 for Tris vs 2’3’-cGAMP comparisons and |*t*| ≤ 0.112, *p* ≥ 0.985 for Tris vs c-di-GMP) but not in mutant flies (|*t*| ≤ 1.547, *p* ≥ 0.222) and a significant increase in survival in control but not in mutant flies (*z* = 2.404, *p* = 0.032 and *z* = - 0.433, *p* = 0.665 for *dSTING^Control^* and *dSTING^Rxn^* flies for the pairwise comparisons between Tris and 2’3’-cGAMP after a Cox proportional hazards model). Relative viral loads at different time points of control flies after co-injection of Tris, 2’3’-cGAMP or c-di-GMP with CrPV (**c**), VSV (**d**), FHV(**e**) or KV (**f**). Co-injection with 2’3’-cGAMP, but not c-di-GMP led to a significantly reduced accumulation of all tested viruses (|*t*| ≥ 2.276, *p* ≤ 0.049 and |*t*| ≤ 1.769, *p* ≥ 0.148 for all pairwise comparisons of Tris vs 2’3’-cGAMP or c-di-GMP at the different days). Data are from two or three independent experiments. Expression levels are shown relative to the housekeeping gene RpL32 and are normalized by experiment.

### 2’3’-cGAMP act independently of the siRNA response and autophagy, but depends upon the NF-κB protein Relish for its antiviral role

To identify the mechanism by which 2’3’-cGAMP exerts its antiviral activity, we first analyzed the effect of CDNs on DCV and VSV infection in *AGO2* null mutant flies. We observed a reduced accumulation of viral RNAs when 2’3’-cGAMP was co-injected with the viruses in both mutant and control flies, revealing that the antiviral function of the CDN does not depend on this key component of the antiviral siRNA pathway (Fig.4a,b). Similarly, 2’3’-cGAMP significantly reduced viral RNA accumulation in *Atg7* null mutant flies, ruling out an involvement of this essential component of the canonical autophagy pathway (Fig.4c). By contrast, the protective effect of 2’3’-cGAMP was completely abolished in *Relish* mutant flies (Fig.4d). Altogether, these results reveal that 2’3’-cGAMP triggers a dSTING-Relish-dependent antiviral transcriptional response, rather than RNA interference or autophagy.

**Figure 4.**
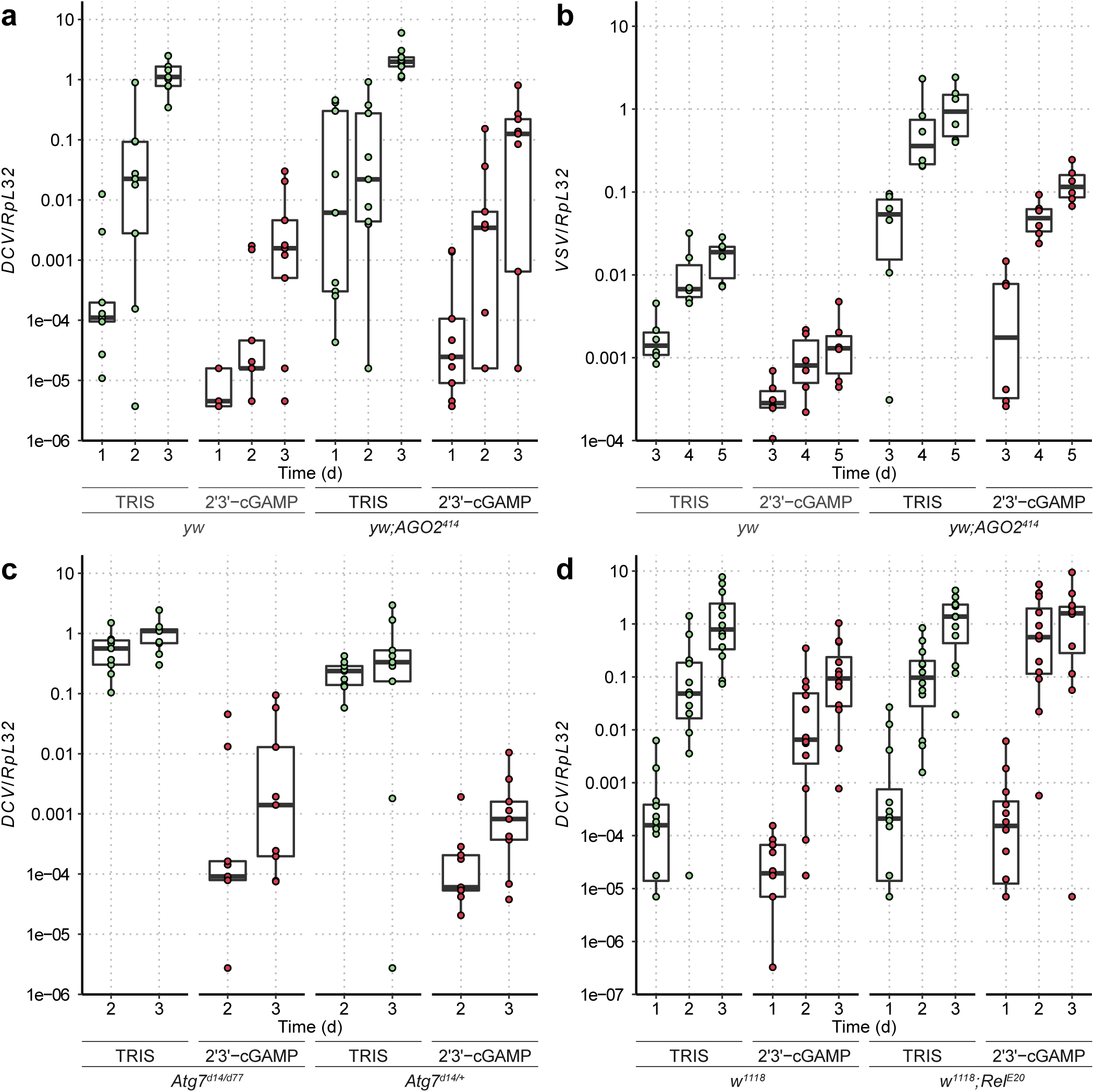
2’3’-cGAMP induced antiviral protection is dependent on Relish, but not on Atg7 or AGO2. Viral RNA loads at different time points after co-injection of Tris or 2’3’-cGAMP with DCV or VSV in flies mutant for the RNAi pathway (*yw;Ago2^414^*, **a,b**), autophagy (*Atg7^d14/d77^*, **c**), Relish (*w^1118^;Rel^E20^*, **d**), or in control flies of the same genetic background (*yw*, *Atg7^d14^/CG5335^d30^ - Atg7^d14/+^*or *w^1118^*, respectively). Co-injection with 2’3’-cGAMP led to a reduced accumulation of viral RNAs in RNAi or autophagy impaired flies (|*t*| ≥ 2.30, *p* ≤ 0.024) and in their controls (|*t*| ≥ 2.53, *p* ≤ 0.013) but not in Relish mutants (|*t*| ≤ 1.220, *p* ≥ 0.225). Data are from two or four (**d**) independent experiments. Expression levels are shown relative to the housekeeping gene *RpL32* and are normalized by experiment. Triangles indicate points where viral RNA could not be detected: threshold cycles (Cq) values for these points were replaced by the maximum Cq for a virus infected sample + 1.

## DISCUSSION

### CDNs activate antiviral immunity in Drosophila

Our results reveal that three out of the four naturally occurring CDNs that activate mammalian STING can also trigger the dSTING signaling pathway in flies. They raise the question of the mechanism by which 2’3’-cGAMP activates dSTING. Like others, we have not been able to detect binding of 2’3’-cGAMP to purified recombinant dSTING^20^. The recently reported cryoEM structure of full length chicken STING reveals substantial interaction of the ligand binding domain with areas of the transmembrane domains at the N-terminus of the protein^26^. We believe that such interaction may be critical for either ligand binding or stability (or both) of the cytosolic domain of dSTING, a hypothesis supported by the differences between the transmembrane domains of STING in mammals and drosophila (e.g. only three transmembrane helices in flies).

Our work complements the molecular study of Kranzusch and colleagues, who reported binding of CDNs to STING from the sea anemone *N. vectensis*, and supports the hypothesis that the ancestral function of STING in metazoans was to sense CDNs^20^. Bacteria produce a diversity of CDNs and cyclic trinucleotides, some of which could activate dSTING^17, 27^. Of note, bacterial CDNs have two canonical 3’,5’ phosphodiester-linkages, whereas mammalian and *Nematostella* cGAS produce chemically distinct CDNs containing one 2’,5’ phosphodiester bond joining G to A and one canonical 3’,5’-phosphodiester bond joining A to G^15, 20^. While we detected activity of 3’3’-CDNs, namely of 3’3’-cGAMP and of c-di-AMP, the strongest agonist was 2’,3’-cGAMP, suggesting that an enzyme producing this CDN exists in insects. Indeed, Wang and colleagues recently reported the inducible production of cGAMP in the cytosol of *Bombyx mori* cells infected with nucleopolyhedrovirus (NPV)^18^. Thus, the production of CDNs in the response to virus infection appears to be ancient, possibly inherited in early eukaryotes from prokaryotes^16, 28^. A major goal for future study will be the identification and characterization of the cGAS enzyme operating in *Drosophila*.

### Activation of NF-**κ**B is an ancestral function of the dSTING pathway

One major difference between STING in mammals and invertebrates, e.g. *Nematostella* and *Drosophila*, is the lack of the CTT domain that mediates interaction with and activation of the kinase TBK1 and the IRF3 transcription factor^16^. This has led to the hypothesis that invertebrate STING regulates autophagy rather than a transcriptional response. Indeed, STING activates autophagy through a mechanism independent of TBK1 activation and IFN induction in mammals. Furthermore, NvSTING also induces autophagy when it is ectopically expressed in human cells^15^. In *Drosophila*, dSTING-dependent autophagy has been proposed to restrict Zika virus infection in the brain, although autophagy constituents are proviral for Zika and other flaviviruses in mammalian cells^19, 29^. Our results using *ATG7* mutant flies indicate that 2’3’-cGAMP controls viral infection independently from the canonical autophagy pathway. However, we cannot rule out the involvement of an unconventional autophagy pathway. Indeed, LC3 lipidation in response to cGAMP stimulation in human cells does not depend on the ULK kinases or Beclin 1, two essential components of the classical autophagy pathway^15^. In this regard, we note that one of the genes upregulated by cGAMP is *ref(2)P*, the ortholog of the autophagy receptor p62 and a restriction factor for Sigma virus^24^. Even though we cannot completely rule out a contribution of autophagy, our results point to the central role played by the NF-κB transcription factor Relish in the antiviral response triggered by 2’3’-cGAMP. Further analysis will be required to precisely define the contribution of Relish in this response. The dSTING-dependent transcriptional response to cGAMP injection is complex, involving up and down regulation of gene expression occurring in different waves, with early and late responses. However, the presence of consensus binding sites for NF-κB in the *cis*-regulatory regions of ∼75% of the regulated genes, regardless of their kinetics of induction and repression, confirms a major contribution of Relish. In addition, we identified 13 other transcription factors and 2 cytokines (*upd3* and *spz*) in the early and sustained up-regulated genes (Table S3). Among the upregulated transcription factors, *kay* (the Drosophila ortholog of c-Fos), *Ets21C* and *FoxK* were previously implicated in immune, inflammatory or stress responses in Drosophila^30–32^. Importantly, the *cis*-regulatory regions of the differentially expressed genes were enriched for binding sites for the mentioned transcription factors and STAT92E, the sole Drosophila STAT ortholog (Fig3d, Table S4). These different transcriptional regulators may coordinate the kinetics of the response and induction of different sets of genes in the context of bacteria and virus infection.

### A broad antiviral induced response in *Drosophila*

We observed a striking antiviral activity of 2’3’-cGAMP against a broad range of viruses with DNA or RNA genomes. This contrasts with previous studies that reported virus-specific induced responses, leading to the idea that RNA interference is the only pathway acting on the broad range of viruses infecting invertebrates, which are devoid of interferons. Importantly, we showed that the antiviral effect of 2’3’-cGAMP does not require AGO2, a key component of the antiviral RNAi pathway in flies, even though this gene is upregulated by the CDN. Thus, besides RNAi, an induced antiviral response involving dSTING contributes to host defense against a range of viruses in *Drosophila*. Furthermore, the induction of AGO2 by CDNs suggest a crosstalk between the two pathways where activation of dSTING potentiates the siRNA response. Intriguingly, our previous results pointed to a specific contribution of the dSTING-IKKβ-Relish pathway in resistance to DCV and CrPV, although a significant but smaller effect was visible also for VSV^5^. This apparent discrepancy could be explained by differences between viruses in the induction of the pathway based on their tissue tropisms, the type of virus-associated molecular pattern produced or the existence of escape strategies, all of which may be bypassed by systemic injection of 2’3’-cGAMP.

A number of previous studies reported strong transcriptional responses to virus infection in insects^2, 21, 25, 33–35^, but also *C. elegans*^36^, oysters^37^ and shrimps^38^. Analysis of the transcriptional response to viral infections *in vivo* is complicated by the fact that (i) cell infections are unsynchronized; (ii) host cells are modified through hijacking of cellular functions by viruses; and (iii) many viruses trigger cell lysis and tissue damage, making it complicated to discern the immune response from the non-specific response to stress. Consequently, the transcriptome of virus-infected flies only provides a blurred image of the induced antiviral response^2, 21, 25, 34, 35, 39^. Identification of an agonist of dSTING bypasses the need for the use of viruses and provides a much clearer picture of the modifications of the drosophila transcriptome associated with induction of antiviral immunity. In particular, our data suggest that 2’3’-cGAMP triggers the expression of cytokines (e.g. Spaetzle, upd3) that amplify the response and trigger expression of antiviral effectors (e.g. Nazo, vir-1). The tools are now at hands to characterize the induced mechanisms controlling viruses in insects, which may reveal original targets for antiviral therapy.

## ACKNOWLEDGEMENTS

This work was supported by the ERANET Infect-ERA program (ANR-14-IFEC-0005), the Agence Nationale de la Recherche (ANR-17-CE15-0014), the Investissement d’Avenir Programs (ANR-10-LABX-0036 and ANR-11-EQPX-0022), the Chinese National Overseas Expertise Introduction Center for Discipline Innovation” (Project “111” (D18010)), CNRS and INSERM. Further support was provided to by the Novo Nordisk foundation grant NNF17OC0028184.

## AUTHOR CONTRIBUTIONS

H.C., A.H., B.S., J.S., A.L., J.H., and J.H. performed experiments; N.M. performed bioinformatics analysis; D.C. and T.P. provided critical materials; H.C., A.H., J.T.M., R.H., N.M. and J.L.I. designed the experiments and analyzed the data; H.C., R.H., N.M. and J.L.I. wrote the manuscript.

## COMPETING FINANCIAL INTERESTS

The authors declare no competing financial interests.

## FIGURE LEGENDS

**Figure S1**

DCV infection induces a dSTING dependent transcriptional response in *D. melanogaster.* **a** *dSTING^Rxn^* mutant flies were generated by imprecise excision of the P-element P{EPgy2}Sting^EY06491^. Precise excision of the transposon generated control flies in the same genetic background. **b-e** Relative gene expression at different days (d) post-injection of Tris or DCV for *dSTING* (**b**) and *srg1-3* (**c-e**) in control flies (*dSTING^Control^*), *dSTING^Rxn^* mutant flies and *dSTING^Rxn^* mutant flies containing a genomic *dSTING* rescue transgene (*dSTING^Rescue^*).

Expression of dSTING was significanly lower in *dSTING^Rxn^* mutant (*t* ≤ −7.189, *p* ≤ 0.001 in all pairwise comparisons between control and *dSTING^Rxn^* in the different timepoints) and identical to control levels in rescue flies (|*t*| ≤ 2.044, *p* ≥ 0.142 in all pairwise comparisons between control and *dSTING^Rescue^* in the different timepoints). STING is induced by DCV infection in *dSTING^Rxn^* mutant flies (|*t*| >= 3.632, *p* ≤ 0.001 for all pairwise comparisons between Tris and DCV injected *dSTING^Rxn^*) and reaches levels close to wild type three days post infection (|*t*|= −2.466, *p* = 0.065 for the comparison between DCV injected *dSTING^Rxn^*and Tris injected control flies). Induction of *srg1* was lower at three dpi in *dSTING^Rxn^* mutants, induction of *srg2* was lower at 3 dpi (*t* = 0.6252, *p* = 0.002) and levels of *srg3* were similar in Tris and DCV infected *dSTING^Rxn^* mutants 2- and 3-dpi (|*t*| ≤ 1.268, *p* >= 0.446) and always lower than in control flies (> 4.85 fold, |*t*| >= 5.568, *p* > 0.001). All these genes were induced by DCV infection in control or *dSTING^Rescue^* flies two or three dpi (|*t*|>= 2.520, |*p*| ≤ 0.037), except for srg3 in control flies 2 dpi (*t* = 1.393, *p* = 0.373) (**f**) Expression levels of *Hsp26*, a dSTING-independent virus induced gene. Induction of *Hsp26* by DCV was identical in control and *dSTING^Rxn^* (|*t*| < 0.842, *p* > 0.405, comparison of differences in *Hsp26* levels between Tris and DCV injected flies at 2 or 3dpi) Data are from two independent experiments. Expression levels are shown relative to the housekeeping gene *RpL32* and are normalized by experiment.

**Figure S2**

Venn diagram of the (**a**) up- and (**b**) down-regulated genes between 2’3’-cGAMP and Tris injected *dSTING^Control^*flies at the different timepoints (6, 12 and 24h) after injection or on average across all timepoints.

**Table S1**

Differentially expressed genes between Tris and c-di-GMP injected *dSTING^Control^* flies at 6, 12 and 24 hours post-injection. Columns represent Ensembl gene ID (**gene_id**) and symbol (**gene_symbol**), mean normalized counts after Tris (**TRIS_**) or c-di-GMP injection (**c-di-GMP_**) at the different timepoints (**_06**,**_12** or **_24**), together with estimated log_2_(fold-change) (**lfc_**) and Benjamini-Hochberg corrected *p* values for the comparison between c-di-GMP and Tris injected flies at each individual timepoint and on average across all timepoints (**_AVG**).

**Table S2**

Differentially expressed genes between Tris and 2’3’-cGAMP injected *dSTING^Control^* flies at 6, 12 and 24 hours post-injection. Columns represent Ensembl gene ID (**gene_id**) and symbol (**gene_symbol**), temporal expression category (**category**) mean normalized counts after Tris (**TRIS_**) or 2’3’-cGAMP injection (**cGAMP_**) at the different timepoints (**_06**,**_12** or **_24**), together with estimated log_2_(fold-change) (**lfc_**) and Benjamini-Hochberg corrected *p* values for the comparison between c-di-GMP and Tris injected flies at each individual timepoint and on average across all timepoints (**_AVG**).

**Table S3**

Differentially expressed transcription factors or cytokines between Tris and 2’3’-cGAMP injected *dSTING^Control^* flies at 6, 12 and 24 hours post-injection. Columns headings are as in Table S2, but include the transcription factor family/sub-family (**Family**).

**Table S4**

Differentially expressed genes between Tris and 2’3’-cGAMP injected *dSTING^Control^* flies with regulatory sequences enriched for the differentially expressed transcription factors, or for transcription factors of the same family/sub-family. Columns headings are as in Table S3, and include the high confidence transcription factor calls predicted by Rcistarget.

## METHOD DETAILS

### Drosophila strains

Fly stocks were raised on standard cornmeal agar medium at 25°C. All fly lines used in this study were *Wolbachia* free. *w^1118^*, *dSTING^Control^*, *dSTING^Rxn^*, *yellow (y) white (w) DD1, yw;AGO2^414^, Atg7^d14^/Cyo-GFP*, *Atg7^d77^/Cyo-GFP* and *CG5335^d30^/Cyo-GFP* stocks have been described previously^40^*. Relish^E20^* flies isogenized to the DrosDel *w^1118^* isogenic background were a gift from Dr. Luis Teixeira^41^.

The genomic rescue of wild-type dSTING was established by PhiC31 mediated transgenesis. The fosmid FlyFos015653^42^ was injected into the *y^1^ w^1118^*; *PBac{y[+]-attP-9A}VK00027* (BDSC#9744) line and introgressed into a *dSTING^Rxn^*mutant background by standard genetic crossing techniques. Transgenesis and initial recombinant fly selection was done by the company BestGene (Chino Hills, CA, USA).

### Virus infection

Viral stocks were prepared in 10 mM Tris-HCl, pH 7.5. Infections were performed with 3–5 days old adult flies by intrathoracic injection (Nanoject II apparatus, Drummond Scientific) with 4.6 nL of DCV solution (500 PFU/fly). Injection of the same volume of 10 mM Tris-HCl, pH 7.5, was used as a negative control.

### CDNs injection with or without viruses

The CDNs (Invivogen) were dissolved in 10 mM Tris pH 7.5 to a concentration of 0.9 mg/mL. 3–5 days old adult flies were CDN stimulated. For CDN injection, each fly was injected with 69 nL of CDN solution or 10 mM Tris pH 7.5 (negative control) by intrathoracic injection using a Nanoject II apparatus. For CDNs and viruses coinjection, 30 μL 0.9 mg/mL CDNs were mixed with 2 μL viruses (DCV 5PFU/4.6 nL, CRPV 5PFU/4.6 nL, VSV 5000PFU/4.6 nL, FHV 500PFU/4.6 nL and KV). Each fly was injected with 69 nL of CDNs or 10 mM Tris pH 7.5 plus virus mixture by intrathoracic injection using a Nanoject II apparatus (Drummond Scientific) and injected flies were collected at indicated time points and homogenized for RNA extraction and RT-qPCR.

### RNA extraction and qRT-PCR

Total RNA from collected flies was extracted using a Trizol Reagent RT bromoanisole solution (MRC), according to the manufacturer’s instructions. 1 μg total RNA was reverse transcribed using an iScript™ gDNA clear cDNA synthesis Kit (Biorad), according to the manufacturer’s instructions. The DNase and RNA reaction mixture was incubated for 5 min at 25°C to remove genomic DNA and then the reaction was stopped by heating at 75°C for 5min. Then reverse transcription mix was added to DNase-treated RNA template and cDNA was synthesized in the following PCR program: Step 1: 25°C for 5 min, step 2: 46°C for 20 min, step 3: 95°C for 1 min. cDNA was used for quantitative real time PCR (QRT-PCR), using iQ^TM^ Custom SYBR Green Supermix Kit (Biorad) according to the manufacturer’s instructions, on a CFX384 Touch™ Real-Time PCR platform (Bio-Rad). Primers used for real-time PCR are listed in Supplementary Table 1. Normalization was performed relative to the housekeeping gene *RpL32*.

### RNA-Sequencing of *D. melanogaster* injected with CDNs

Forty male flies of *dSTING^Control^* and *dSTING^Rxn^* were injected with 69 nL/fly of either 10 mM Tris (pH 7.5), c-di-GMP (1mg/mL) or 2,3-cGAMP (1 mg/mL) by intrathoracic injection (Nanoject II apparatus), in three independent experiments. Injected flies were collected at 6-, 12- and 24-hours post injection. Total RNA was isolated from injected flies using TRIzol™ Reagent (Invitrogen), according to the manufacturer’s protocol. RNA quantity and purity were assessed using a Dw-K5500 spectrophotometer (Drawell) and Agilent 2200 TapeStation (Agilent). rRNA was removed using Epicentre Ribo-Zero rRNA Removal Kit (Illumina), and RNA was converted to cDNA. Prepared cDNA was used for Illumina sequencing library preparation using NEBNext® Ultra™ Directional RNA Library Prep Kit for Illumina (NEB), following the manufacturer’s instructions. Briefly, DNA fragments were end repaired to generate blunt ends with 5′phosphatase and 3′hydroxyls, before adapters ligation, PCR amplification and cleanup. Average fragment length was 300-bp. Purity of the libraries was evaluated using an Agilent 2200 TapeStation. Libraries were used for cluster generation in situ on an HiSeq paired-end flow cell using the Rapid mode cluster generation system, followed by massively parallel sequencing (2×150 bp) on an HiSeq X Ten. Library construction, high throughput sequencing, adapter removal and initial quality control and trimming were done by the company Ribobio (Guangzhou, China) The data set for CDNs injected flies were submitted to the GEO database (Gene Expression Omnibus: http://www.ncbi.nlm.nih.gov/geo/), with the accession numbers (to be communicated).

### Transcriptome analysis

After quality trimming and adapter removal using Trimmomatic, reads were mapped using STAR v2.5.3^43^ to the Drosophila genome and annotation (ENSEMBL BDGP6.22). Reads mapping to the sense strand of the transcripts were counted with featureCounts v1.6.2^44^, using the Drosophila annotation files, allowing mapping to multiple genes. Differential gene expression of transcripts present in ≥20% of the libraries with at least 5 reads across all libraries was done using the *deseq* function of the “DESeq2” (v1.20) package^45^. Variance was estimated using the local fitting method. Read counts and normalized read counts are shown in GEO datasets (to be communicated). Transcripts with log2 difference in expression ≥ 1.5 and Benjamini & Hochberg corrected *p*-value < 0.05 were considered differentially expressed.

### Clustering of temporal expression profiles

All differentially expressed genes between Tris and 2’3’-cGAMP injected WT flies at any time point or on average across all time points were clustered in temporal expression categories by partitioning around medoids (PAM) clustering using the *pam* function in the “cluster” (v2.1.0) package. The optimal number of clusters for either up- or down-regulated genes was determined using the gap statistic method, as implemented in the *fviz_nbclust* function of the “factoextra” (v1.0.5) package, using default parameters (100 bootstrapped replications, 10 maximum allowed clusters). Gene expression clusters were visualized using the *Heatmap* function of the “ComplexHeatmap” (v2.0.0) package and *ggplot* of the “ggplot2” (v3.2.1) package.

### Ontology analysis

Differentially expressed genes between Tris and 2’3’-cGAMP injected WT flies in each temporal expression category were tested for enrichment relative to all genes passing the expression cutoff in any gene ontology type (Molecular Function, Cellular Compartment, Biological Process), using the “Generic GO subset” of gene ontology terms (downloaded from http://current.geneontology.org/ontology/subsets/index.html on 10/10/2019). Gene ontology enrichment analysis was done using the *enricher* function of “clusterProfiler” package (v3.1.12), using default parameters (Benjamini & Hochberg corrected *p-*value cutoff of 0.05).

### Transcription Factor Enrichment Analysis

Enrichment of transcription factor binding sites (TFBS) in the regulatory regions of the differentially expressed genes was done using the *cisTarget* function of the “RcisTarget” package (v1.4.0)^46^. The database “dm6-5kb-upstream-full-tx-11species.mc8nr” database was used, which includes the rankings for conserved TFBS in the non-coding regions 5 kb upstream of the transcription start site and in introns of all annotated genes in the D. melanogaster genome (r6.02). Gene symbols were updated to the r6.04 annotation when necessary. Transcription factor family assignment was done according to Flybase (FB2019_05).

### Statistical analysis

For quantification of viral RNA loads and target gene expression, log transformed ratios were compared using linear regression models using the *lm* function of base R. Survival curves were analysed by Cox regression using the *coxph* function in the “survival” (v2.44-1.1) package. Depending on the experiment, independent variables included genotype, virus injection, CDN injection and time post injection and all interactions between them. Experiment was included as an independent variable in all tests. Multiple comparisons between the groups of interest and controls were done using the *emmeans* function of the “emmeans” (v1.4.1) package, using Dunnett’s method for *p* values correction. Data were analysed using R (v3.4.2) and *ggplot* was used for plotting.

