## Supplementary figures and images for "2’3’-cGAMP triggers a STING and NF-κB dependent broad antiviral response in Drosophila"

### Figure S1

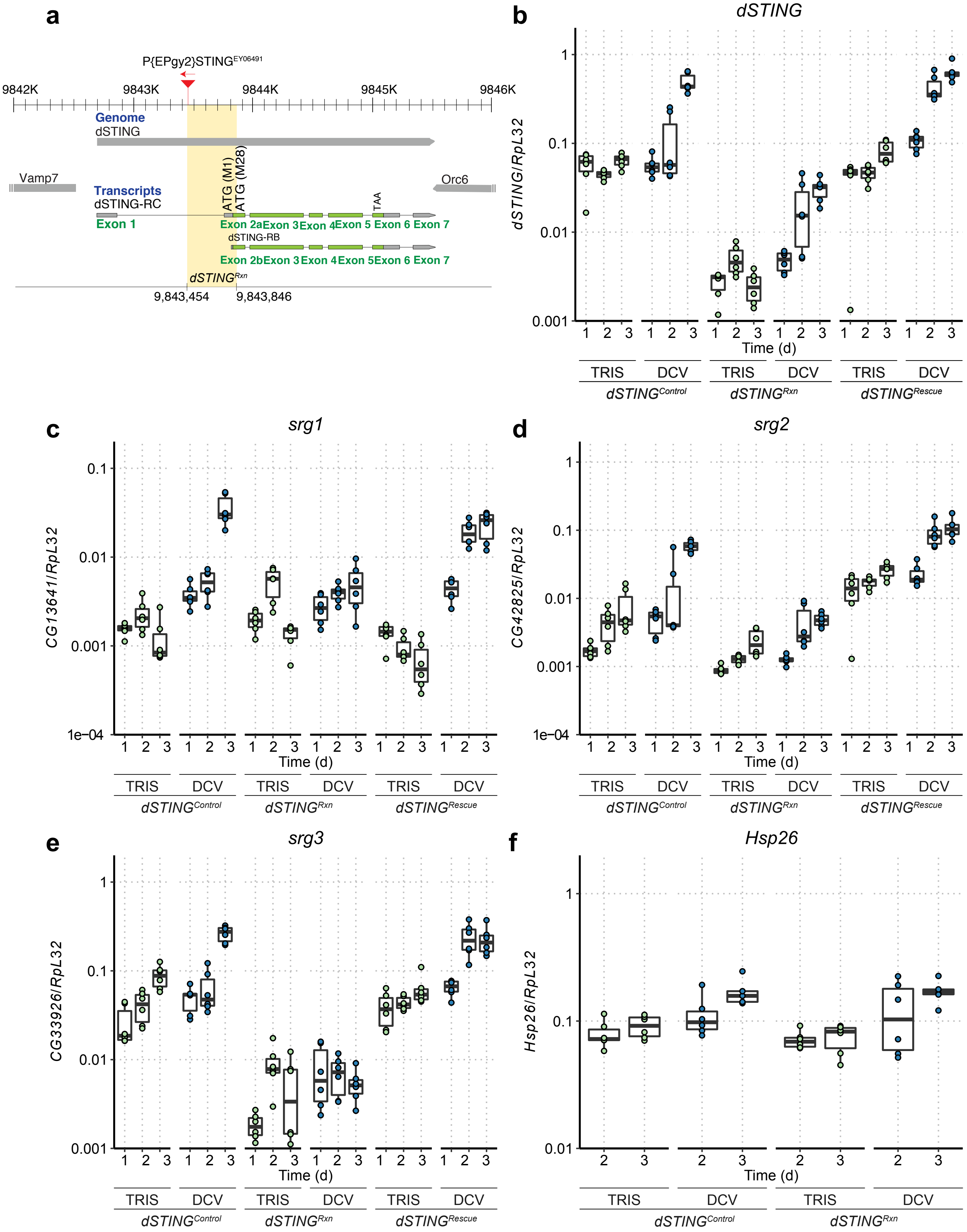


Figure S1

Hua Cai *et al*

### Figure S2

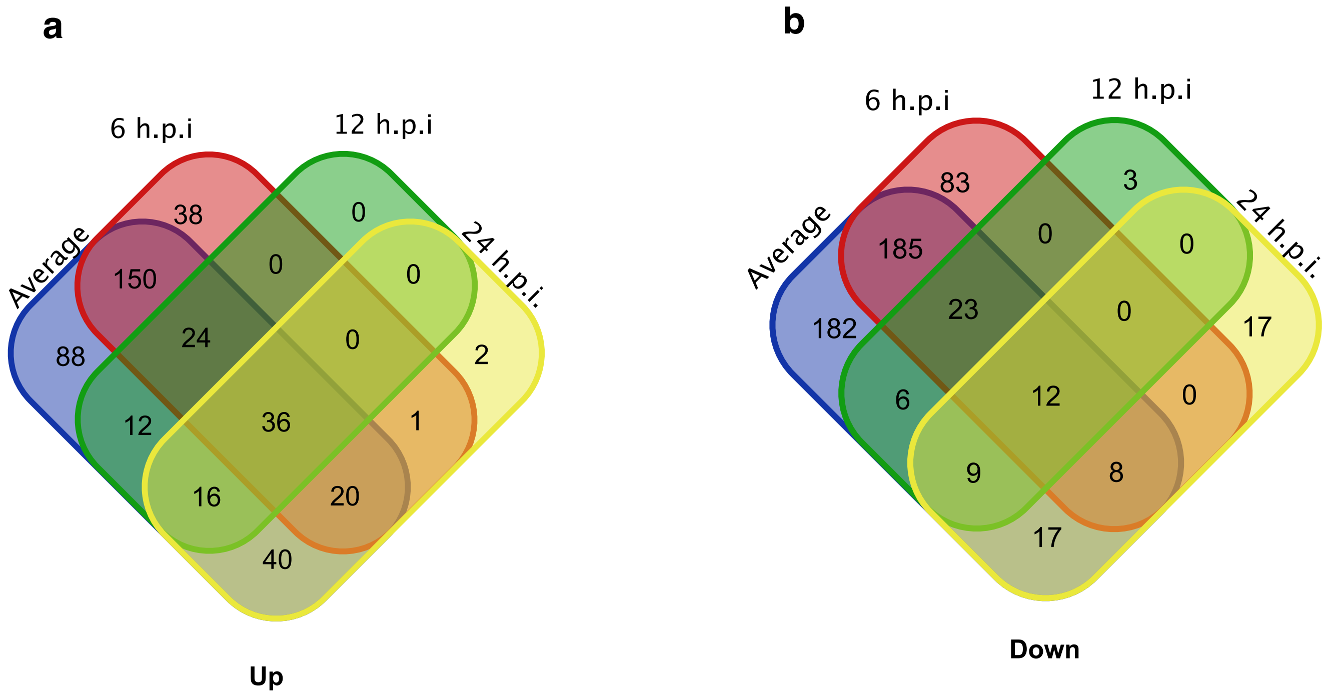


Figure S2

Hua Cai *et al*
